# Functional heterogeneity of cell populations increases robustness of pacemaker function in a numerical model of the sinoatrial node tissue

**DOI:** 10.1101/2021.12.27.474308

**Authors:** Alexander V. Maltsev, Michael D. Stern, Edward G Lakatta, Victor A. Maltsev

## Abstract

Each heartbeat is initiated by specialized pacemaker cells operating within the sinoatrial node (SAN). While individual cells within SAN tissue exhibit substantial heterogeneity of their electrophysiological parameters and Ca cycling, the role of this heterogeneity for cardiac pacemaker function remains mainly unknown. Here we investigated the problem numerically in a 25×25 square grid of coupled-clock Maltsev-Lakatta cell models and tested the hypothesis that functional heterogeneity of cell populations increases robustness of SAN function. The tissue models were populated by cells with different degree of heterogeneity of the two key model parameters of the coupled-clock system, maximum L-type Ca current conductance (*g_CaL_*) and sarcoplasmic reticulum Ca pumping rate (*P_up_*). Our simulations showed that in the areas of *P_up_-g_CaL_* parametric space at the edge of the system stability where action potential (AP) firing was absent or dysrhythmic in tissues populated by identical cells, rhythmic AP generation was rescued in tissues populated by cells with uniformly random distributions of *g_CaL_* or *P_up_* (but keeping the same average values). This effect to increase robust AP generation was synergistic with respect to heterogeneity in both *g_CaL_* and *P_up_* and was further strengthened by clustering of cells with higher *g_CaL_* or *P_up_*. The effect of functional heterogeneity was not due to a simple summation of activity of intrinsically firing cells naturally present in SAN; rather AP firing cells locally and critically interacted with non-firing/dormant cells. When firing cells prevailed, they recruited many dormant cells to fire, strongly enhancing overall SAN function. And vice versa, prevailing dormant cells suppressed AP firing in cells with intrinsic automaticity and halted SAN automaticity.

## INTRODUCTION

The sinoatrial node (SAN) of the right atrium is the heart’s primary pacemaker, providing both robust and flexible automatic electrical depolarizations that capture the surrounding atrial myocardium and drive the heartbeat. These electrical depolarizations are caused by concurrent operation of pacemaker cells residing in SAN tissue. These cells are electrical oscillators that spontaneously generate rhythmic changes of their membrane potential (*V_m_*), producing relatively periodic spontaneous action potentials (APs) (reviews (Mangoni and Nargeot, 2008;Maltsev et al., 2014)). The SAN was discovered in 1907 (Keith and Flack, 1907). However, despite enormous amount of experimental data, numerical modeling, and plausible theories, the SAN operation remains still mysterious after all these years (Weiss and Qu, 2020).

The enigma of SAN function is that the SAN is an extremely complex heterogeneous tissue regarding individual cell properties and their connections within the tissue (Boyett et al., 2000). Furthermore, at the cellular level, individual SAN cells operate via a complex coupled-clock paradigm (Maltsev and Lakatta, 2009;Lakatta et al., 2010), i.e. they possess not only an electrical oscillator within their cell surface membranes, but also an intracellular oscillator, sarcoplasmic reticulum (SR) that cycles Ca in-and-out via SR Ca pump (SERCA2a) and release channels, ryanodine receptors (RyRs). Ca releases via RyRs interact with Na/Ca exchanger and L-type Ca current (*I_CaL_*), and together they drive diastolic depolarization via a positive feedback mechanism (dubbed ignition (Lyashkov et al., 2018)). The synchronization of molecular actions, in turn, is driven by enhanced enzymatic activity of Ca-activated adenylate cyclases (AC1 and AC8) (Mattick et al., 2007;Younes et al., 2008) and protein kinases (PKA and CaMKII) (Vinogradova and Lakatta, 2009;Lakatta et al., 2010) that modulate respective ion channel properties and accelerate Ca pumping.

The expression of Ca cycling proteins, and membrane ion channels vary substantially among individual SAN cells (Honjo et al., 1996;Honjo et al., 1999;Lei et al., 2001;Musa et al., 2002;Monfredi et al., 2017). For example, *I_CaL_* density varies by an order of magnitude (Monfredi et al., 2017) (Fig. 1A, red squares). This results in a large degree of functional heterogeneity among individual cells, ranging from frequently and rhythmic AP firing to infrequently and dysrhythmic firing, to nonfiring cells (recently dubbed “dormant” cells) (Kim et al., 2018;Tsutsui et al., 2018;Tsutsui et al., 2021). The SAN cell population is also extremely heterogeneous with respect to the AP firing rate increase in response to β-adrenergic stimulation (Kim et al., 2021).

**Fig. 1.**
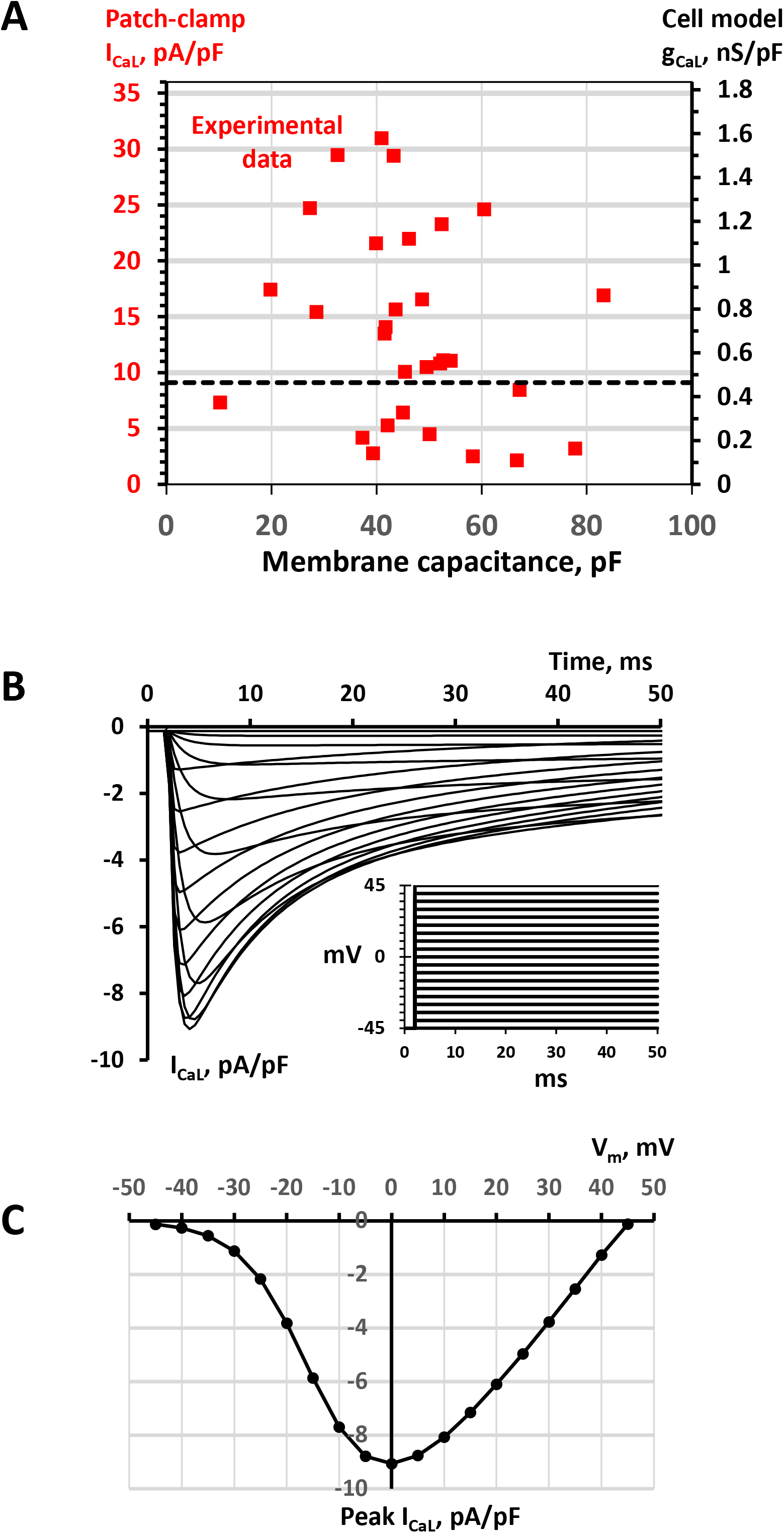
SAN cell model simulations to match model parameter *g_CaL_* to *I_CaL_* density measured experimentally. A: A double Y scale plot of experimental data points of *I_CaL_* densities measured by whole-cell patch clamp in a large population of SAN cells (red squares, left Y axis, replotted from (Monfredi et al., 2018)) and respective model *g_CaL_* parameter values (right Y axis). B: Traces of *I_CaL_* / *V_m_* relationship simulated in Maltsev-Lakatta single cell model with the basal state value of *g_CaL_* =0.464 nS/pF using voltage clamp protocol (inset) and conditions similar to those in experimental study of (Monfredi et al., 2018). The family of *I_CaL_* traces was simulated by applying a series of square voltage pulses from a holding potential of −45 mV with a step of 5 mV up to 45 mV (inset). Before each stimulation, *I_CaL_* activation and inactivation gating variables *f_L_* and *d_L_* were set to their steady-state values. To closely simulate experimental wholecell patch clamp conditions in (Monfredi et al., 2018), 5 mM EGTA (equilibrated to 100 nM of free Ca) was added to Ca formulations of cell submembrane space and cytoplasm. C: Respective *I_CaL_* / *V_m_* relationship calculated for peak *I_CaL_*. The *I_CaL_* peak of 9.068 pA/pF at 0 mV was set as a bridge between experimental data and model *g_CaL_* =0.464 nS/pF (black dash line in panel A).

At the SAN tissue level, it was initially thought that a “master” pacemaker cell or a leading pacemaker center dictates the excitation rate and rhythm of other SAN pacemaker cells (Sano et al., 1978;Bleeker et al., 1980). Subsequent studies suggested that individual cells mutually entrain each other to fire APs with a common period (dubbed “democratic” process) (Jalife, 1984;Michaels et al., 1987). More recent high-resolution imaging of intact SAN tissue at a single cell level, however, demonstrated that while majority of SAN cells indeed fire synchronously with a common period, many cell fire “at will”, i.e. at various rates and irregularly, or remain silent, generating only local Ca releases that do not advance into AP-induced Ca transients (Bychkov et al., 2020;Fenske et al., 2020). Thus, how structural and functional heterogeneities within and among SAN cells give rise to synchronized AP firing at the SAN exits remains a mystery at the frontier of pacemaker research (Clancy and Santana, 2020;Weiss and Qu, 2020).

Numerical modeling is a powerful approach to study complex cell interactions. Here we used a modified 2-dimensional model of SAN tissue developed by Campana (Campana, 2015), featuring faster parallel computing via graphics processing unit (GPU) to test the hypothesis that functional heterogeneity of cells within SAN tissue increases robustness of its pacemaker function. We adapted the model of SAN tissue to include diverse cell populations with respect to maximum SR Ca pumping rate (*P_up_*) and *I_CaL_* conductance (*g_CaL_*) as observed experimentally (Honjo et al., 1996;Monfredi et al., 2017)(Fig. 1A, Table 1). We found that SAN tissue populated by heterogenous cells could generate rhythmic AP firing within the areas of *P_up_* - *g_CaL_* parametric space where SAN populated by identical cells lacked automaticity or generated arrhythmias. The effect was synergistic with respect to *P_up_* and *I_CaL_* and the robustness of heterogeneous SAN tissue could be further increased by clustering cells with similar properties.

**Table 1.** Parameter values in each model simulation scenario (column #) and major results of our simulations during time interval from 5 s to 7.5 s after simulation onset, the percentage of AP firing cells in tissue, the percentage of cells firing APs in isolation, and AP cycle length average with its standard deviation (SD) calculated for all AP firing cells populating SAN model tissue model (25 x 25 square grid) in each scenario. N.D. means not defined in scenarios 4 and 7, in which SAN tissue lacked automaticity.

| # | Distr. | $P_{up}$,<br>min,<br>mM/<br>s | $P_{up}$,<br>max,<br>mM/s | $P_{up}$<br>mean,<br>mM/s | $g_{CaL}$,<br>min,<br>nS/pF | $g_{CaL}$,<br>max,<br>nS/pF | $g_{CaL}$<br>mean,<br>nS/pF | %<br>firing<br>cells,<br>tissue | % firing<br>cells,<br>separate | Cycle<br>length<br>average,<br>ms | Cycle<br>length<br>SD,<br>ms |
| --- | --- | --- | --- | --- | --- | --- | --- | --- | --- | --- | --- |
| 1 | random | 7 |  |  | 0.14 | 0.54 | 0.34 | 64.64 | 46 | 405.91 | 11.72 |
| 2 | random | 5 |  |  | 0.17 | 0.57 | 0.37 | 70.4 | 46 | 429.43 | 9.4 |
| 3 | random | 3 |  |  | 0.21 | 0.61 | 0.41 | 83.2 | 47.5 | 450.3 | 11.55 |
| 4 | random | 2 | 12 | 7 | 0.34 |  |  | 0 | 39 | N.D. | N.D. |
| 5 | random | 0 | 10 | 5 | 0.37 |  |  | 97.92 | 40 | 460.99 | 57.09 |
| 6 | random | 0 | 6 | 3 | 0.41 |  |  | 100 | 40 | 506.91 | 6.27 |
| 7 | random | 7 |  |  | 0.3 | 0.38 | 0.34 | 0 | 30 | N.D. | N.D. |
| 8 | random | 2 | 12 | 7 | 0.3 | 0.38 | 0.34 | 69.12 | 42 | 417.2 | 36.38 |
| 9 | $g_{CaL}$<br>cluster | 7 | | | 0.3 | 0.38 | 0.34 | 60.8 | 30 | 457.28 | 8.39 |
| 10 | $P_{up}$<br>cluster | 2 | 12 | 7 | 0.34 | | | 66.4 | 39 | 368.21 | 33.13 |

## MATERIALS AND METHODS

### Single cell model

Our SAN tissue model is comprised of single cell models developed by Maltsev and Lakatta in 2009 (Maltsev and Lakatta, 2009). Each model encompasses a system of 29 first order differential equations (see state variables *y_1_-y_29_* in Table 2). This was the first SAN cell numerical model featuring coupled operation of Ca and membrane clocks. The model predicted the contribution of spontaneous Ca release during diastolic depolarization via activation of inward Na/Ca exchanger current that explained numerous experimental data. The computer code for the original model can be freely downloaded in CellML format (maltsev_2009_paper.cellml) at http://models.cellml.org/workspace/maltsev_2009 and run using the Cellular Open Resource (COR) software developed at the University of Oxford by Garny et al. (Garny et al., 2009) (for recent development of the COR see http://www.opencor.ws/).

**Table 2.**
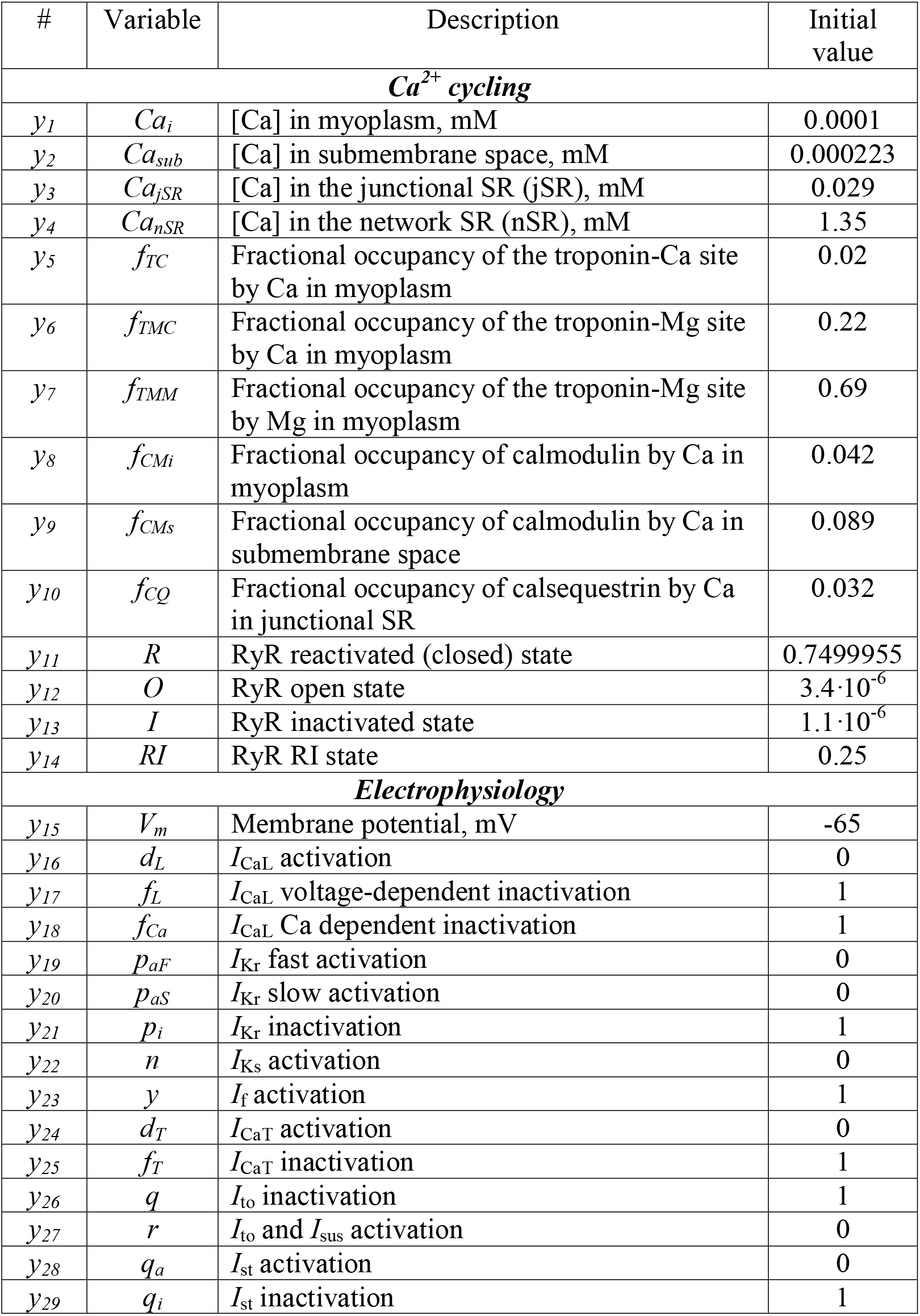
State variables *y_1_-y_29_* with their descriptions and initial values assigned to all cells in our SAN tissue model simulations.

| # | Variable | Description | Initial value |
| --- | --- | --- | --- |
| ***Ca<sup>2+</sup> cycling*** |  |  |  |
| $y_1$ | $Ca_i$ | [Ca] in myoplasm, mM | 0.0001 |
| $y_2$ | $Ca_{sub}$ | [Ca] in submembrane space, mM | 0.000223 |
| $y_3$ | $Ca_{jSR}$ | [Ca] in the junctional SR (jSR), mM | 0.029 |
| $y_4$ | $Ca_{nSR}$ | [Ca] in the network SR (nSR), mM | 1.35 |
| $y_5$ | $f_{TC}$ | Fractional occupancy of the troponin-Ca site by Ca in myoplasm | 0.02 |
| $y_6$ | $f_{TMC}$ | Fractional occupancy of the troponin-Mg site by Ca in myoplasm | 0.22 |
| $y_7$ | $f_{TMM}$ | Fractional occupancy of the troponin-Mg site by Mg in myoplasm | 0.69 |
| $y_8$ | $f_{CMi}$ | Fractional occupancy of calmodulin by Ca in myoplasm | 0.042 |
| $y_9$ | $f_{CMs}$ | Fractional occupancy of calmodulin by Ca in submembrane space | 0.089 |
| $y_{10}$ | $f_{CQ}$ | Fractional occupancy of calsequestrin by Ca in junctional SR | 0.032 |
| $y_{11}$ | $R$ | RyR reactivated (closed) state | 0.7499955 |
| $y_{12}$ | $O$ | RyR open state | $3.4 \cdot 10^{-6}$ |
| $y_{13}$ | $I$ | RyR inactivated state | $1.1 \cdot 10^{-6}$ |
| $y_{14}$ | $RI$ | RyR RI state | 0.25 |
| ***Electrophysiology*** |  |  |  |
| $y_{15}$ | $V_m$ | Membrane potential, mV | -65 |
| $y_{16}$ | $d_L$ | $I_{CaL}$ activation | 0 |
| $y_{17}$ | $f_L$ | $I_{CaL}$ voltage-dependent inactivation | 1 |
| $y_{18}$ | $f_{Ca}$ | $I_{CaL}$ Ca dependent inactivation | 1 |
| $y_{19}$ | $p_{aF}$ | $I_{Kr}$ fast activation | 0 |
| $y_{20}$ | $p_{aS}$ | $I_{Kr}$ slow activation | 0 |
| $y_{21}$ | $p_i$ | $I_{Kr}$ inactivation | 1 |
| $y_{22}$ | $n$ | $I_{Ks}$ activation | 0 |
| $y_{23}$ | $y$ | $I_f$ activation | 1 |
| $y_{24}$ | $d_T$ | $I_{CaT}$ activation | 0 |
| $y_{25}$ | $f_T$ | $I_{CaT}$ inactivation | 1 |
| $y_{26}$ | $q$ | $I_{to}$ inactivation | 1 |
| $y_{27}$ | $r$ | $I_{to}$ and $I_{sus}$ activation | 0 |
| $y_{28}$ | $q_a$ | $I_{st}$ activation | 0 |
| $y_{29}$ | $q_i$ | $I_{st}$ inactivation | 1 |

Since 2009, the model has been tested, modified and used in numerous applications (Maltsev and Lakatta, 2010;Kurata et al., 2012;Severi et al., 2012;Maltsev and Lakatta, 2013;Campana, 2015;Sirenko et al., 2016;Kim et al., 2018;Lyashkov et al., 2018;Yang et al., 2021). According to the coupled-clock theory, a coupled system of Ca and surface membrane “clocks” can provide the robustness and flexibility required for normal pacemaker cell function. A simple numerical release-pumping-delay Ca oscillator is capable of generating any frequency between 1.3 and 6.1 Hz, but cannot generate high-amplitude signals without the assistance of membrane clocks (Maltsev and Lakatta, 2009). The coupled-clock system utilizes the greater flexibility of the SR Ca clock while simultaneously accounting for the large Ca oscillation amplitudes fueled via sarcolemmal Ca influx via *I_CaL_*. A modern interpretation of the coupled-clock function is that Ca clock operates via a criticality mechanism, whereas the membrane clock operates as a limited cycle oscillator, comprising a perfect synergistic pacemaker cell system (Weiss and Qu, 2020).

### Multi-cellular model of SAN

The present study examined performance of SAN tissue that was modelled as a square grid of 25 by 25 of Maltsev-Lakatta cell models (Maltsev and Lakatta, 2009), with each cell being connected to its four neighbors (except those on the border). Our model was an adapted version of the tissue model originally developed by Campana (Campana, 2015). The final formulation to compute voltage in a 2-dimensional network of SAN cells was as follows (equation 2.7 in (Campana, 2015)):

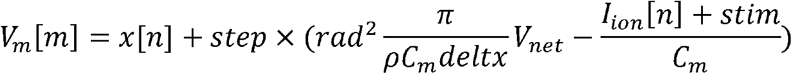

Where *V_m_[m]* is *V_m_* of the cell with index *m* within the cellular network represented by a vector (1..number of cells); *x* is the state variable array; for each time step, *m* and *n* denote the same cell, only in two different vectors and *x[n]* represents the value assumed by *V_m_[m]* at the previous time step; *V_net_* is the sum of voltages from the four neighboring cells (right, left, up, and down): *V_net_=V_net_R_+V_net_L_+V_net_U_+V_net_D_* (Fig. 2); *step* is the model integration time step of 0.005 ms; *rad* is the cell radius of 4 μm; *ρ* is the intracellular resistivity of 10^4^ MΩm; *C_m_* is the electrical membrane capacitance of 32 pF; *deltx* is the cell length of 70 μm; *I_ion_* is the sum of all ion currents; *stim* is stimulus current (*stim*=0 in our study).

**Fig. 2.**
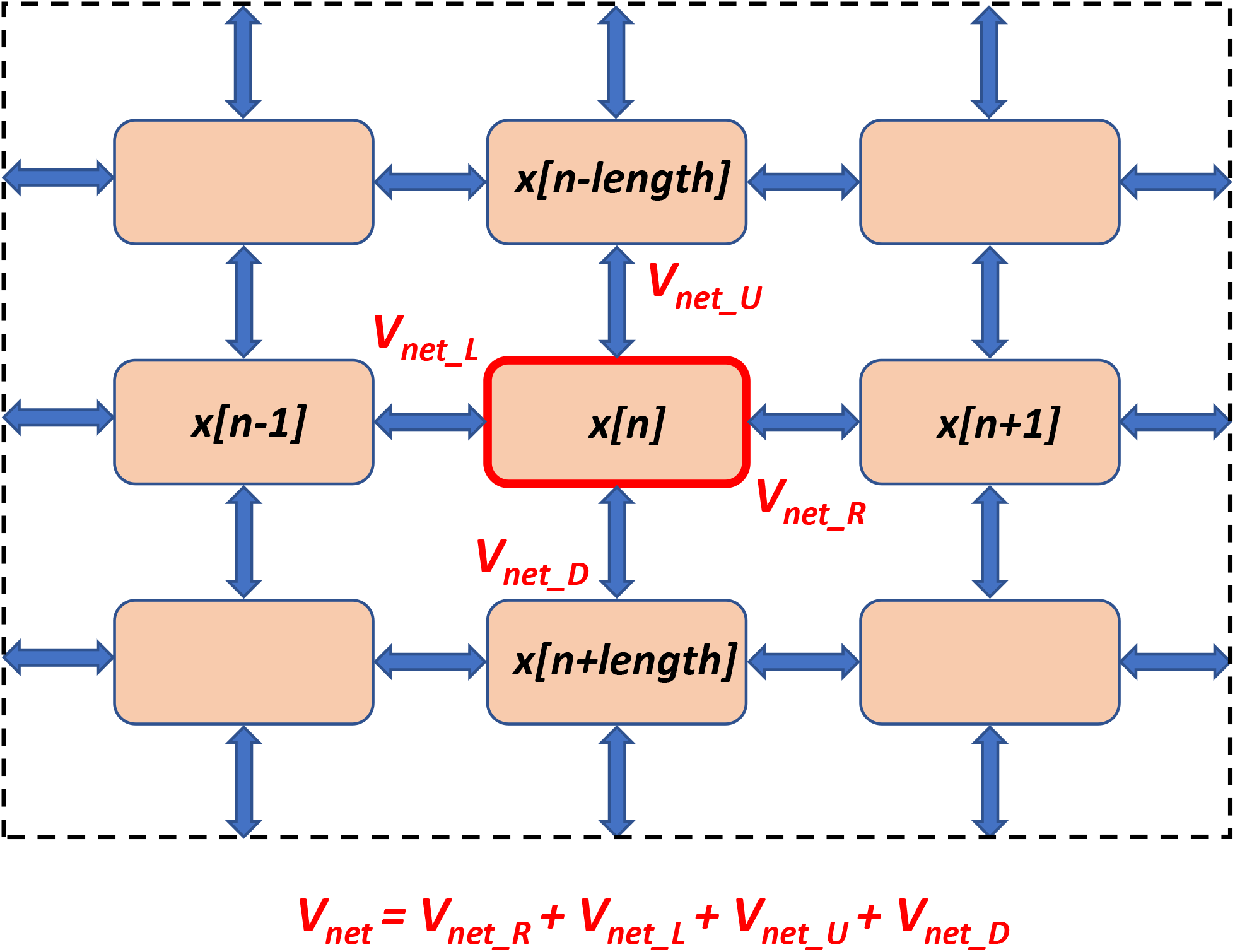
A schematic illustration of SAN tissue structure and calculation of the network voltage (*V_net_*) for a cell with index *n* in Campana model (Campana, 2015). Pink squares denote individual SAN cells (Maltsev-Lakatta models) located at the nodes of a square grid. The cells interact via intercellular conductances placed uniformly at the grid’s edges (shown by blue double-headed arrows). *V_net_* is the sum of voltages from the four neighboring cells (right, left, up, and down). Shown is only a 3×3 element the network; the tissue continues beyond the dash lines to form the entire 25×25 cell network used in majority of our simulations.

### Simulation of functional heterogeneity in SAN tissue models

Tissue heterogeneity was introduced in the model by varying *P_up_* and *g_CaL_*, the two key parameters of the coupled-clock system (Maltsev and Lakatta, 2009). *P_up_* determines the rate at which Ca is pumped into the SR by SERCA2a to reach a threshold of Ca load required for activation of Ca release, i.e. a central part in the timing mechanism of the coupled-clock system (Maltsev and Lakatta, 2009;Imtiaz et al., 2010;Vinogradova et al., 2010;Stern et al., 2014;Maltsev et al., 2017). Our cell model approximated Ca uptake flux (*j*_up_) to the network SR compartment as a function of cytoplasmic [Ca] (*Ca_i_*) as suggested originally by Luo and Rudy (Luo and Rudy, 1994) in ventricular cells and subsequently used in SAN cell models, e.g. (Kurata et al., 2002;Maltsev and Lakatta, 2009;Severi et al., 2012):

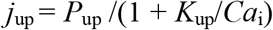

where *K*_up_ = 0.6·10^-3^ mM is *Ca_i_* for a half-maximal Ca uptake to the network SR. Thus, in our simulations of heterogeneous SAN tissue we only varied *P_up_*, with *K*_up_ remaining constant. *I_CaL_* as a function of *V_m_*, submembrane [Ca] (*Ca_sub_*), and time (*t*) was approximated as follows (for more details, see (Maltsev and Lakatta, 2009)):

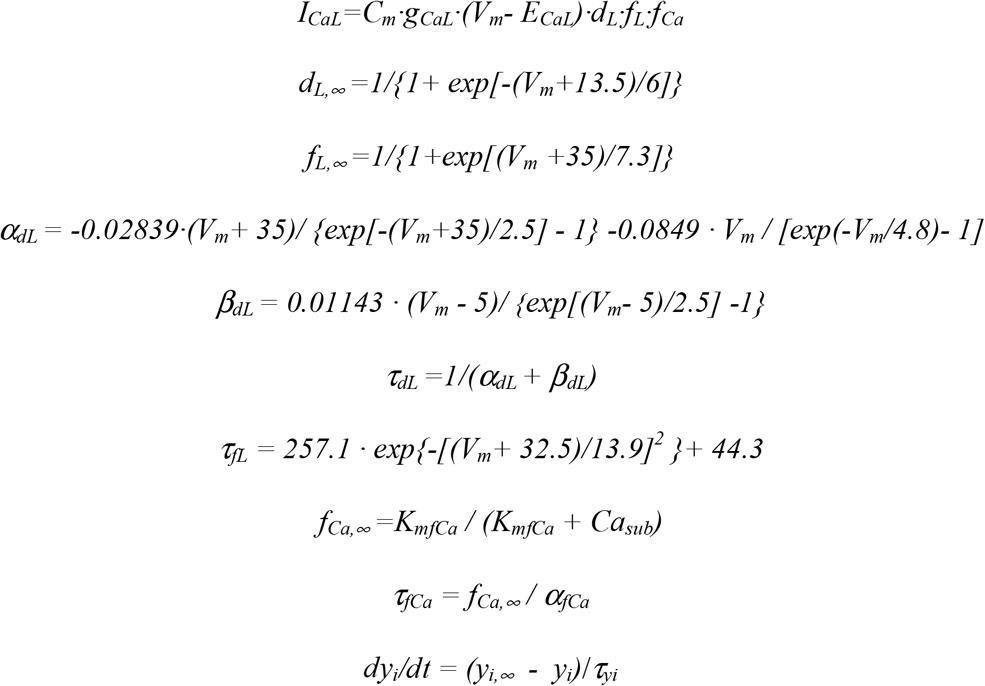

where *y_i_* = *d*_L_, *f*_L_, *f*_Ca_ are respective channel gating variables of Hodgkin-Huxley type (state variables *y_16_, y_17_, y_18_* in Table 2) with steady-state probabilities *d_L,∞_, f_L,∞_, f_Ca,∞_* and gating time constants *τ_dL_, τ_fL_, τ_fCa_*. *E_CaL_* = 45 mV is an apparent reversal potential of *I_CaL_* and *C_m_* = 32 pF is electric capacitance of the cell membrane.

In simulations of heterogeneous SAN tissue, we varied only *g_CaL_*, maintaining all other parameters unchanged. In other terms, only number of functional channels in each cell was varied, but channel gating kinetics remained the same in all cells. We directly linked *g_CaL_* to *I_CaL_* density measured experimentally by whole patch-clamp technique under volage clamp conditions (Fig. 1, see details in Results). Once we bridged our model to experimental data, we used a *P_up_* - *g_CaL_* bifurcation diagram previously reported in single cell models (Maltsev and Lakatta, 2009) to guide our selection of *P_up_* and *g_CaL_* distributions for cell populations that fell into parameter-dependent areas of rhythmic firing, chaotic firing, or no firing, bordering at a bifurcation line (split yellow line in Fig. 3A). Since our study was focused on the system robustness, we examined the tissue operation in the areas of no firing or chaotic firing (dubbed non-firing zone and chaotic firing zone, respectively) in which individual cells lack normal automaticity but normal pacemaker function is possible in heterogeneous tissue models. The parameters for cell populations were set via a uniform random distribution within specified ranges (Table 1). For some cell populations, we also tested the effect of cell clustering within the SAN grid.

**Fig 3.**
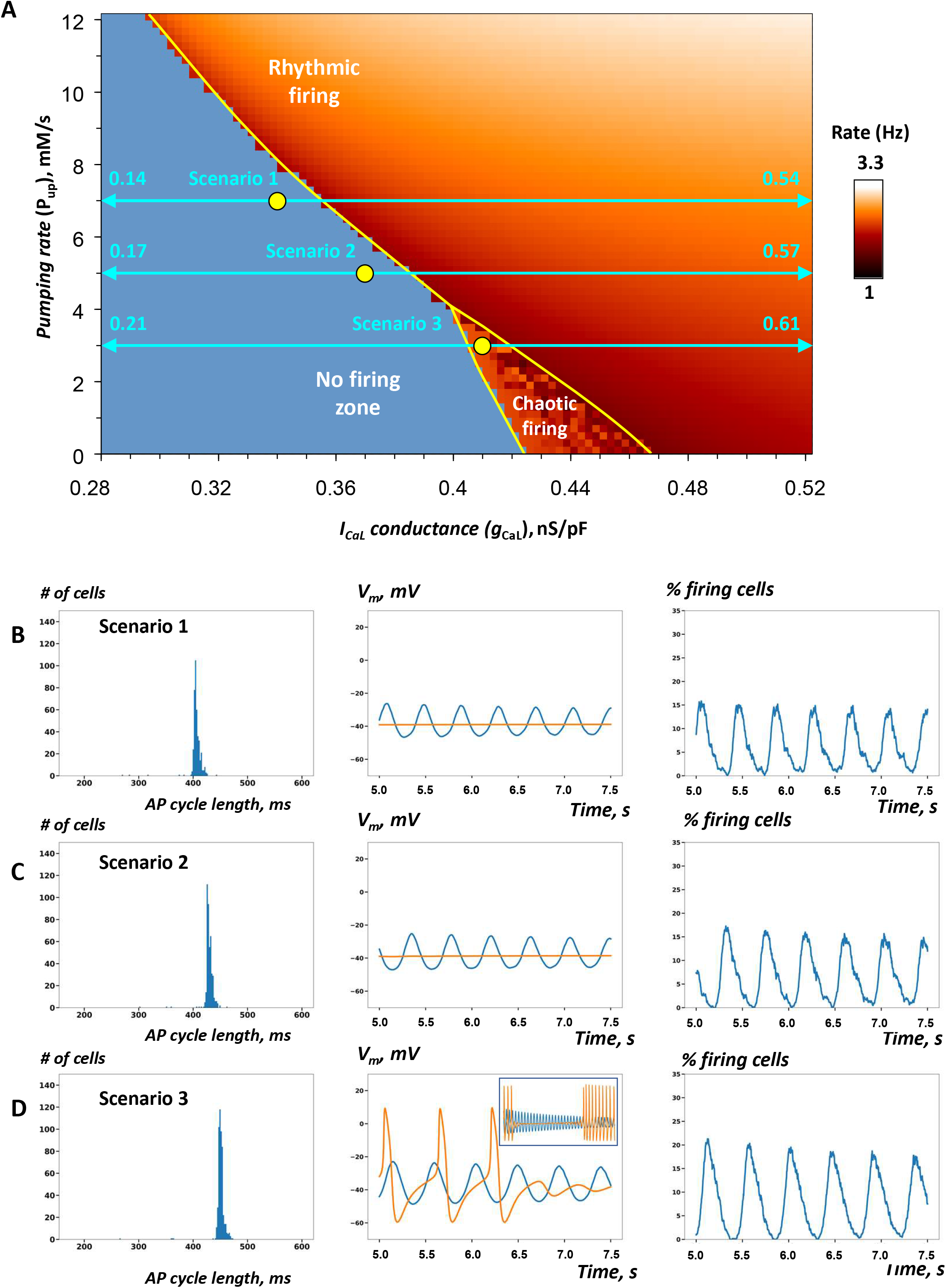
Heterogeneity in *g_CaL_* increases robustness of AP firing in SAN tissue models close to the edge of stability. A: *P_up_- g_CaL_* bifurcation diagram previously reported for single SAN cell model (modified from (Maltsev and Lakatta, 2009)) shows the parametric space for rhythmic AP firing, no firing, and chaotic firing (white labels). Yellow circles show coordinates for fixed *P_up_* and *g_CaL_* mean values (all in the non-firing zone) in cell populations used in our simulations of SAN tissue function in 3 specific scenarios 1-3 (Table 1) close to the bifurcation line (yellow line). Double-headed aqua arrows show the *g_CaL_* ranges for each scenario (Note: the ranges extend beyond the diagram, with the labels showing the extensions). All three tissue models fired rhythmic APs (Movies 1-3). B-D: Each panel shows the results of simulations in each scenario: histograms of AP cycle length distribution, space-average *V_m_* vs. time, and the percentage of firing cells (*V_m_*>0) vs. time (from 5s to 7.5 s). In scenario 3, we extended the simulation run to 25 s. Inset in panel D (middle) shows average *V_m_* from 5 s to 25 s. Red traces show space-average *V_m_* vs. time for the control SAN tissue model populated by identical cells with average *P_up_* and *g_CaL_* values of the respective heterogeneous model.

### Computation, program code, data analysis and visualization

Our simulations of SAN tissue function were performed using a new computational approach suggested by Campana (Campana, 2015) based on CUDA technology (Compute Unified Device Architecture). A major advantage of this approach is its parallel processing via GPU that is critical to perform simulations of hundreds or even thousands cell models within SAN tissue within a reasonable time.

The model was originally written in CUDA C programming language for a GPU that featured 64 CUDA cores and 768MB RAM (Nvidia Quadro FX 1800) (Campana, 2015). Aside from various code optimizations, we adapted the code to modern, high-performance GPUs such as TITAN RTX graphics card (NVIDIA corp. CA, USA) featuring 4608 CUDA cores and 24 GB RAM used in the present study. One specific important improvement was that we arranged CUDA blocks in a grid, rather than in a one-dimensional array in the original code. Furthermore, random numbers were properly initialized in the main function of Central Processing Unit and copied to GPU memory to generate random values for *g_CaL_* and *P_up_* for each cell. The option for clustering random numbers in the center of the grid was also implemented in the main function.

All simulations began with the same initial conditions, near to the maximum diastolic potential (Table 2). In nearly all simulations, the system reached a steady pattern of AP firing (or no firing) in about 5 s. Thus, our standard simulations lasted 7.5 s, and we examined the system behaviors in the last 2.5 s. Some simulations were run longer, for 25 s, when the system required more time to reach its steady AP firing pattern. In all model simulation scenarios, we compared capabilities of spontaneous AP firing by SAN tissue models populated by heterogenous cells vs. control models populated by identical cells whose *g_CaL_* and *P_up_* were assigned to their respective mean values in heterogenous tissue models. The data analysis and visualizations (including movies of *V_m_* in the tissue) were performed using in-house written programs in Python 3.10.1 (Python Software Foundation, www.python.org) and Delphi 10.4. (Embarcadero, Austin, TX).

## RESULTS

### Matching the experimentally measured *I_CaL_* densities and the model parameter *g_CaL_*

To bridge realistic *I_CaL_* densities (measured in pA/pF by patch clamp (Monfredi et al., 2018), Fig. 1A) and the model parameter, *g_CaL_* (in nS/pF), we performed voltage-clamp simulations of our single cell model (Fig. 1B) with the voltage step protocol (see inset) similar to that used in experimental studies (Honjo et al., 1996;Monfredi et al., 2017). The *I_CaL_* - *V_m_* relationship for the peak values of the simulated *I_CaL_* traces (Fig. 1C) revealed a maximum peak *I_CaL_* current of 9.068 pA/pF at 0 mV with a basal state conductance of 0.464 nS/pA, bridging experimental and model data via black dash line in the dual Y scale plot in Fig. 1A. In our tissue model scenarios involving a large variety of *g_CaL_* values in individual cell models, we always referred to Fig. 1A to make sure that *g_CaL_* remained within the respective range *I_CaL_* densities measured experimentally.

### SAN models with heterogeneous *g_CaL_* close to the bifurcation line

First, we tested the hypothesis that reported heterogeneity in *I_CaL_* density among SAN cells (i.e. *g_CaL_* in the model terms) adds robustness to AP firing within SAN tissue. We constructed scenarios in which the mean *g_CaL_* values were in the non-firing zone along the bifurcation line (yellow circles in Fig. 3A), but with a substantial spread of individual *g_CaL_* values (horizontal double-headed aqua arrows). Three scenarios with different *g_CaL_* distributions were tested to ensure robustness of the simulation results. Our specific choice of *g_CaL_* range in each scenario is given in Table 1 (scenarios 1-3). The respective simulation results are presented in Fig. 3B-D, Table 1, and visualized in Movies 1-3. The percentage of AP firing cells within the time interval from 5 to 7.5 s after simulation onset was the most important parameter, by which SAN tissue performance was evaluated (columns “% firing cells” tissue vs. separate in Table 1).

In all three scenarios, rhythmic AP firing was indeed observed in our simulations of SAN heterogeneous models (Fig.3B-D, blue lines), whereas all models populated with identical cells failed to generate APs (Fig.3B-D, red lines in middle panels). In scenarios 1 and 2, impulses originating within multiple areas in the grid spread in a wave-like fashion to other areas involving the majority of cells (Movies 1 and 2). In scenario 3, rhythmic wave-like AP propagation recruited almost all cells (83.2%), despite mean values of *P_up_* and *g_CaL_* fell into the chaotic firing zone (Movie 3). For scenario 3, we extended our simulation to 25 s to ensure that the control model populated with identical cells indeed generated chaotic firing (Fig. 3D, middle panel inset), rather than just ceased firing after 7.5 s in our shorter standard simulation run. Thus, scenario 3 explicitly shows that adding heterogeneity to *g_CaL_* can have **antiarrhythmic** effect: dysrhythmic (chaotic) firing in SAN tissue model populated by identical cells is shifted to rhythmic firing in the heterogeneous SAN model.

Histograms of AP firing length distribution calculated for all three scenarios (Fig.3B-D, left panels) exhibited sharp peaks between 400 and 500 ms with a relatively small standard deviation of about 10 ms (Table 1). Synchronization of AP firing among cells is also evident from grid-average *V_m_*, oscillation amplitude (about 25 mV, middle panels) and the percentage of firing cells (*V_m_* >0) that oscillated with an amplitude of 15-20% (right panels).

### SAN models with heterogenous *P_up_*

Next, we examined 3 model scenarios (#4, 5, 6 in Table 1) in which *P_up_* was uniformly randomly distributed, but *g_CaL_* was fixed to a certain value in each scenario. The *P_up_* value in a given cell mimics the functional effect of phospholamban phosphorylation to regulate SR Ca pumping via SERCA2a. The mean values of *P_up_* and fixed *g_CaL_* (yellow circles in Fig. 4A) were chosen to move along and close to the bifurcation line, still remaining within the non-firing zone. Here we tested the specific hypothesis that adding heterogeneity to *P_up_* in cell populating SAN tissue increases its robustness to generate rhythmic AP firing.

**Fig 4.**
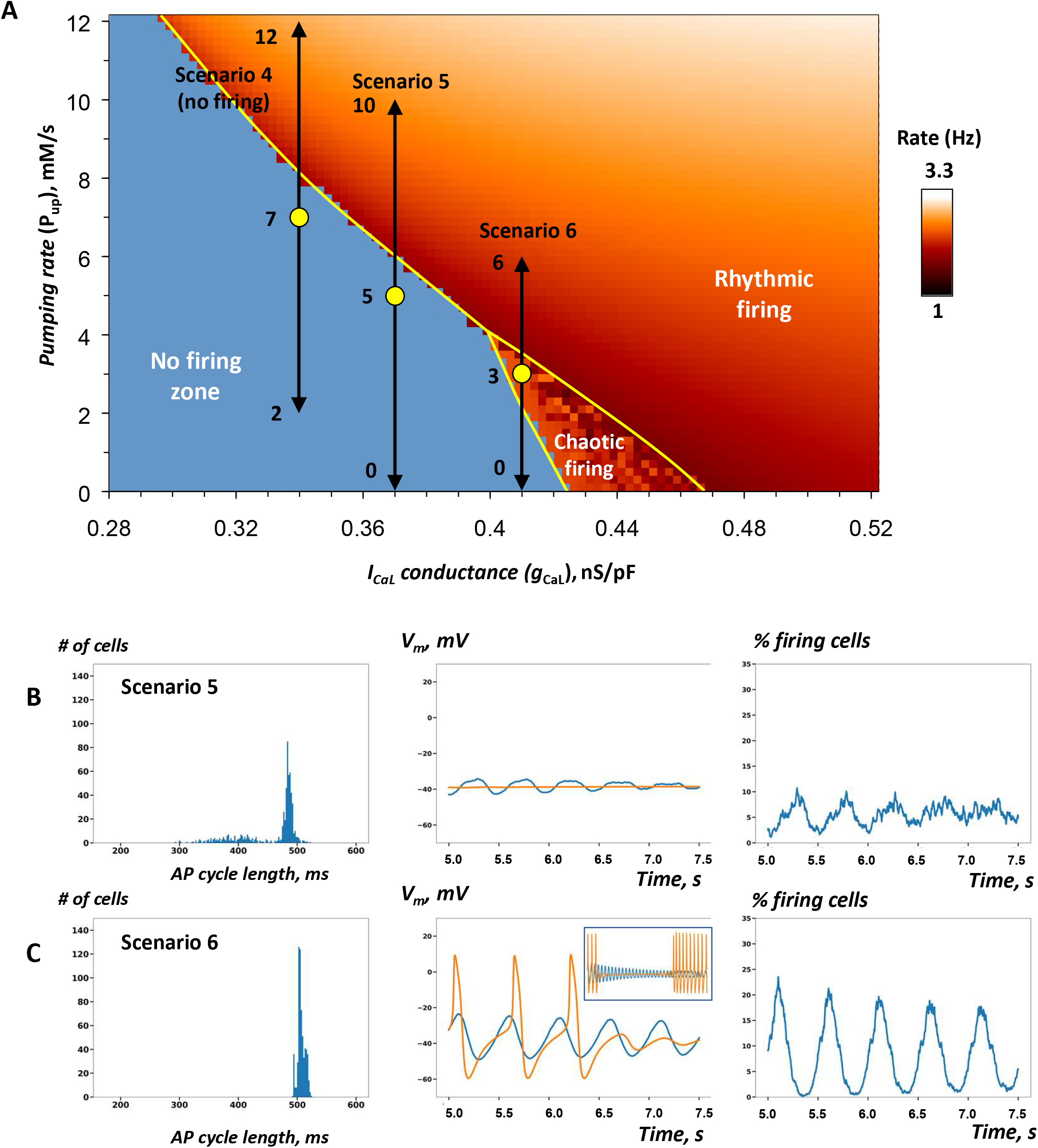
Heterogeneity in *P_up_* increases robustness of AP firing in SAN tissue models close to the edge of stability. A: *P_up_*- *g_CaL_* bifurcation diagram illustrating cell populations in simulations of SAN tissue function in scenarios 4-6 (Table 1) similar to Figs 3A. Yellow circles show coordinates for mean *P_up_* and fixed *g_CaL_* values (in non-firing and the chaotic firing zones). Black double-headed arrows show the exact *P_up_* ranges for each scenario. The tissue model with low *g_CaL_* in scenario 4 failed to generate AP firing despite a higher average *P_up_* value (Movie 4), but scenarios 5 and 6 with higher *g_CaL_* values fired rhythmic AP (Movies 5-6). B-C: the results of simulations in scenarios 5 and 6. In scenario 6, we extended the simulation run to 25 s. Inset in panel C (middle) shows average *V_m_* from 5 s to 25 s.

A substantial cell-to-cell variability in phospholamban phosphorylation is expected based on a substantial variability of both basal AP firing rates and responses to β-adrenergic stimulation among isolated SAN cells (Kim et al., 2021). However, in contrast to *g_CaL_*, there is no single-cell data in the literature to set up cell populations in the model within an exact *P_up_* range. To get an idea of what to expect for physiologically relevant lower limit in *P_up_* variations in individual cells (when testing model stability), one can refer, for example, to how low *P_up_* can descend during the physiological responses to cholinergic receptor stimulation. Within physiologic range of acetylcholine concentrations up to 1 μM, the *P_up_* values can decrease on average from 12 mM/s to as low as 4 mM/s (Fig 4A in (Maltsev and Lakatta, 2010)), indicating that *P_up_* can actually be very flexible in each SAN cell during its normal operation. Thus, we constructed and tested 3 model scenarios with substantial variability of *P_up_* (Fig. 4; Table 1, scenarios 4-6; Movies 4-6). Our model simulations in scenarios 5 and 6 demonstrated that *P_up_* heterogeneity can indeed restore SAN tissue operation within the non-firing zone, thus supporting our hypothesis.

In scenario 4, however, *g_CaL_* (and hence cell excitability) were set so low, that none of the 625 cells in the entire grid continued to generate APs after about 4 s (Movie 4). In scenarios 5 and 6, however, despite the decreasing average of *P_up_* values, the tissue generated APs (Table 1) as *g_CaL_* increased (Table 1). This indicates importance of *g_CaL_* and clock coupling in this mechanism of robustness. Surprisingly and in contrast to simulations involving the *g_CaL_* spread in the previous section, all or almost all (100% or 98%) cells fired rhythmic APs (Table 1). However, the tissue-wide AP firing occurred in a less synchronized manner, especially in scenario 5, manifested by substantial spread in AP cycle lengths among cells (histogram in Fig. 4B, large SD value in Table 1, Movie 5). Furthermore, the percentage of firing cells at a given time was extremely noisy. A large fraction of the grid showed persistent unsynchronized AP firing that was unlike any scenario across *g_CaL_* spread (#1-3) where the mean voltage of the grid exhibited a clear oscillatory pattern. Interestingly, the histogram of AP cycle length in scenario 5 appeared to be bi-modal with two peaks, indicating that cells may self-organize into two subpopulations with different AP cycle lengths.

### Rescuing AP firing requires substantial heterogeneity in *P_up_* or *g_CaL_*

It is important to emphasize that in all above scenarios rescuing AP firing required a substantial spread of *g_CaL_* values (within 0.4 nS/pF) or *P_up_* values. In other terms, in the non-firing zone at each *P_up_* value, the spread of *g_CaL_* must reach a critical value in order to rescue AP firing. While the spread of 0.4 nS/pF is indeed substantial, it was realistic, because assigned *g_CaL_* values remained within experimentally measured range of *I_CaL_* as shown in our bridging dual Y plot in Fig. 1A. We tested several additional scenarios with narrower *g_CaL_* distribution spreads < 0.4 nS/pF, but those non-realistic SAN tissue models were unable to converge to steady AP firing. One such failed tissue model is shown in Fig. 5A (scenario 7, no firing) by aqua double-headed arrow (see also Movie 7).

**Fig 5.**
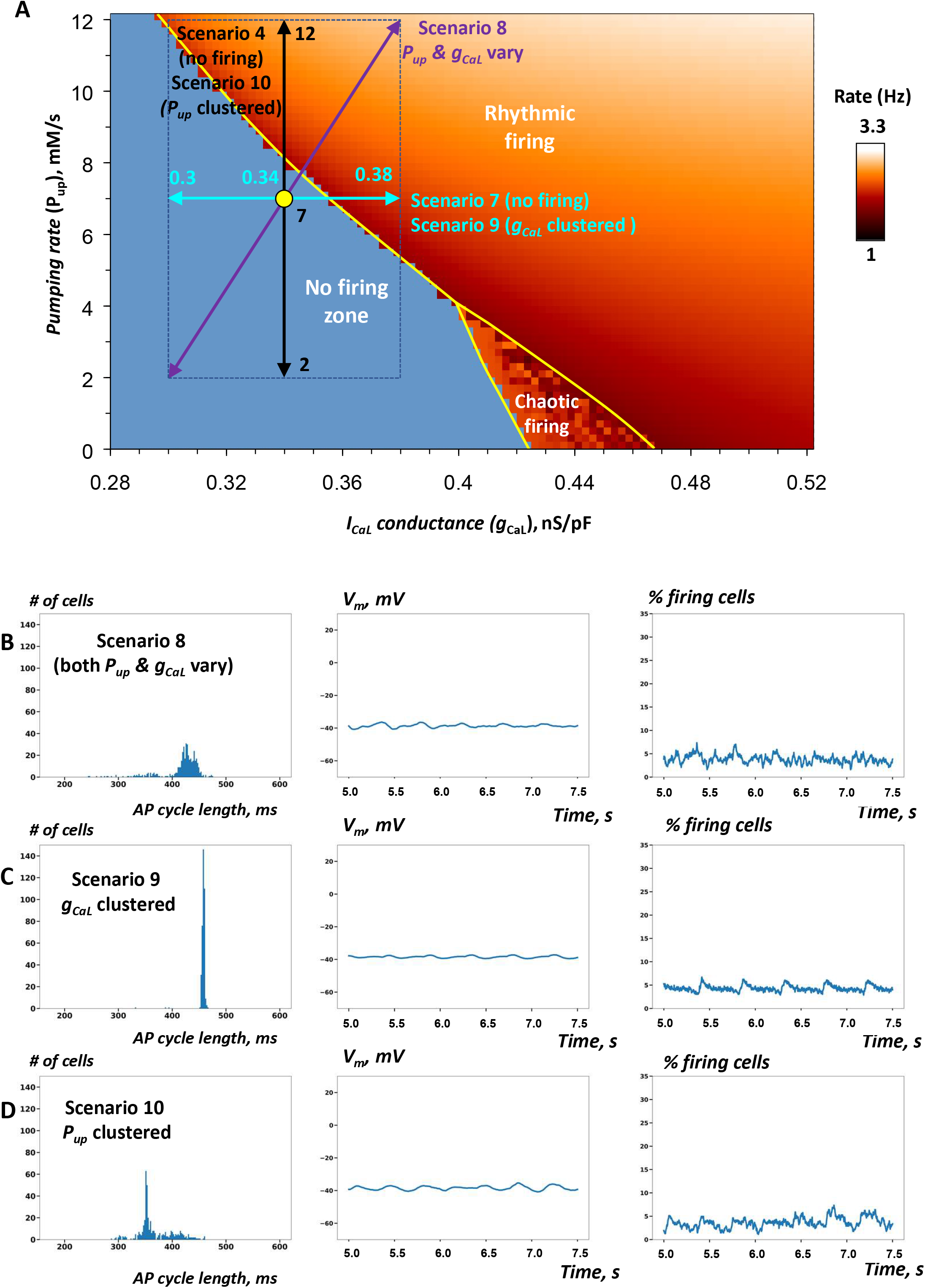
Robustness of SAN operation is increased via combined *P_up_* and *g_CaL_* heterogeneities and their clustering. A: *P_up_-g_CaL_* bifurcation diagram illustrating cell populations in simulations of SAN tissue function in scenarios 7-10 (Table 1) similar to Figs 3A and 4A. Yellow circle shows coordinates for mean *P_up_* and mean *g_CaL_* values (in the non-firing zone). Double-headed arrows show *P_up_* and *g_CaL_* ranges in these scenarios. The tissue models with heterogeneity in either single parameter failed to generate rhythmic APs (Movies 4 and 7). However, when both parameters fluctuated, the tissue model fired rhythmic APs (Movie 8). Furthermore, when either parameter (*g_CaL_* or *P_up_*) was clustered, the tissue also generated APs in scenarios 9 or 10 (Movie 9 or 10), respectively. B-D: the results of simulations in scenarios 8-10.

### Functional synergy of *P_up_* and *g_CaL_* heterogeneities

To gain further insights into SAN tissue operation, we tested the hypothesis that heterogeneity of cell populations with respect to both Ca clock and membrane clock would act synergistically to increase robustness of the pacemaker function. When *P_up_* or *g_CaL_* were distributed separately in scenarios 4 and 7 (described above), the respective SAN tissue models lacked automaticity (Movies 4 and 7). Thus, we simulated and examined an additional special scenario (#8) that included the distributions of both *g_CaL_* and *P_up_* assigned in scenarios 4 and 7 (purple diagonal double-headed arrow in Fig. 5A, Table 1). Our simulations showed that such system with dual parameter distribution generated APs (Fig. 5B, Movie 8), indicating that heterogeneities in *g_CaL_* and *P_up_* act indeed synergistically in rescuing normal system operation within the non-firing zone. Remarkably, the percentage of firing cells in the tissue increased to 69% from 42% for the same cell population when cells are not connected to each other, indicating ongoing recruitment of dormant cells to fire APs when cells were connected. AP firing in this scenario, however, was poorly synchronized among cells with such narrow distribution of *g_CaL_* values around a low mean value of only 0.34 nS/pF.

### *P_up_* and *g_CaL_* clustering is yet another mechanism of robust SAN function

In all previous scenarios *g_CaL_* and *P_up_* were uniformly randomly distributed within a specific range. Experimental studies, however, indicate that the SAN features clusters of cells with different types of activity (Fig. 4 in (Bychkov et al., 2020)). Thus, we tested the hypothesis that the natural cell clustering provides an additional “gear” to enhance robustness of AP firing in SAN tissue. To test this hypothesis, we attempted to rescue automaticity in scenario 7 in which cells had heterogeneous *g_CaL_* within an extremely narrow range from 0.3 to 0.38 nS/pF. Indeed, when cells were uniformly randomly distributed over the tissue, the modelled system of cells lacked automaticity However, the system was able to achieve rhythmic AP firing when the same heterogenous cell population was locally redistributed with higher *g_CaL_* clustering towards the grid center (Fig.5C; Table 1, scenario 9; Movie 9). Finally, we tested the hypothesis that, similar to *g_CaL_*, increased heterogeneity in *P_up_* can also add robustness to the system of interacting cells. Indeed, while the tissue model in scenario 4 lacked automaticity with uniformly random *P_up_* distribution among cells within the tissue (Movie 4), clustering cells with higher *P_up_* towards the tissue center rescued the system automaticity (scenario 10, Fig 5D, Table 1, Movie 10).

## DISCUSSION

### Our new approach to the problem

Heterogeneity of SAN cells has been known and well-documented for a long time (Boyett et al., 2000), but its importance for SAN function remains unclear. Numerical model simulations performed in the present study revealed an interesting and counterintuitive aspect of the problem: While noise and heterogeneity are usually associated with undesirable disturbances or fluctuations, SAN cell heterogeneity actually increases robustness of cardiac pacemaker function. Our work took advantage of well-described properties of single isolated SAN cells in numerous previous experimental and theoretical studies. Specifically, the intrinsic behaviors of individual cells have been characterized by parametric sensitivity analyses (Maltsev and Lakatta, 2009;Kurata et al., 2012) that revealed synergistic contributions of the two key parameters *P_up_* and *g_CaL_* to generate normal automaticity in individual SAN cells as depicted in *P_up_* - *g_CaL_* diagram (panel A in Figs 3–5). We used this diagram to test specific hypotheses in different scenarios in each model simulation (Table 1) and to interpret the simulation results (discussed below). In other terms, on the one hand we investigated how the SAN tissue behaves when it is populated by a large variety of individual cells with known intrinsic properties (i.e. the systems approach). On the other hand, we limited our consideration to only two key model parameters *g_CaL_* and *P_up_* and their variable combinations that, in essence, determine the function of the SAN cell coupled-clock system (i.e. we combined reductionist and systems approaches).

### Comparison with previous numerical studies of SAN tissue

A plethora of numerical models of SAN tissue function have been developed previously, including models in one dimension (Zhang et al., 2001;Huang et al., 2011;Glynn et al., 2014;Li et al., 2018), two dimensions (Michaels et al., 1987;Oren and Clancy, 2010;Campana, 2015;Gratz et al., 2018;Karpaev et al., 2018), and full-scale SAN models in three dimensions (Munoz et al., 2011;Li et al., 2013;Li et al., 2014;Kharche et al., 2017;Mata et al., 2019). These models examined effects of cell-to-cell coupling (Li et al., 2013;Campana, 2015;Gratz et al., 2018;Mata et al., 2019), inter-cellular variability(Campana, 2015;Mata et al., 2019), coupling to other cell types (fibroblasts) (Oren and Clancy, 2010;Karpaev et al., 2018), and normal and abnormal impulse propagation in three dimensions (Munoz et al., 2011;Li et al., 2014;Kharche et al., 2017).

To our knowledge only two numerical studies (Campana, 2015;Mata et al., 2019) have investigated the impact of heterogeneity of cell parameters on SAN tissue function. However, in both studies cell heterogeneity was limited to **only surface membrane clock** parameters and mimicked by a simultaneous randomization of all membrane conductances with respect to their basal values within a fixed percentage range (dubbed sigma). Mata et. al. (Mata et al., 2019) reported time course of synchronization of AP firing vs. sigma. Campana (Campana, 2015) reported that the cell population synchronizes on a rate slightly higher than the one of the isolated cells with the basal values of conductances, but it does not equal the rate of the fastest cell. Our study, in part, confirms this result in scenarios 3 and 6 with chaotic firing in the basal state (Figs 3D and 4C), but provides further insights on specific importance of heterogeneity in *I_CaL_* and *P_up_* for robustness of AP firing in SAN tissue (not studied previously).

Examining Ca signals in addition and together with membrane clock parameters is vital in studies of pacemaker function because each SAN cell operates as a coupled-clock system, i.e. actions of *I_CaL_* and *P_up_* are intertwined in a synergistic manner in numerical models of both isolated cells (Maltsev and Lakatta, 2009;Kurata et al., 2012) and SAN tissue (in the present study, Fig. 5A,B). Among membrane currents, we chose *I_CaL_* for our analysis because it is involved not only in membrane clock function (it generates AP upstroke (Mangoni and Nargeot, 2008), but also in clock coupling by providing influx of Ca (coupled clock’s oscillatory substrate) (Maltsev and Lakatta, 2009;Lakatta et al., 2010) and via positive feedback mechanisms among local Ca releases, Na/Ca exchanger, and *I_CaL_* during diastolic depolarization (Torrente et al., 2016;Lyashkov et al., 2018). Distributions of *I_CaL_* among individual SAN cells has been reported in previous experimental studies (Honjo et al., 1996;Honjo et al., 1999;Lei et al., 2001;Musa et al., 2002;Monfredi et al., 2017) and thus permitted realistic choices for our *I_CaL_* modeling.

Recent theoretical studies have demonstrated that the presence of a Ca clock in addition to a membrane clock protects the SAN from annihilation, associated with sinus node arrest (Li et al., 2018); and that random parameter heterogeneity among oscillators can consistently rescue the system from losing synchrony (Zhang et al., 2021). These prior reports and the results of the present study support a novel SAN structure/function paradigm (resembling neuronal networks) that was proposed based on high-resolution imaging of SAN tissue (Bychkov et al., 2020): in the new paradigm, interactions of Ca signals and APs play a central role in generation of rhythmic cardiac impulses that emanate from the SAN.

### The coupled-clock theory explains, in part, the effect of heterogenous cell populations to enhance robustness of SAN function

Our model simulations performed for SAN tissues operating at the edge of the system stability demonstrated that the robustness of AP firing is enhanced when either *g_CaL_* or *P_up_* varies among the cells (Figs 3 and 4). This result can be explained, in part, on the basis of the coupled-clock theory (Maltsev and Lakatta, 2009). While *P_up_* and *g_CaL_* impact cell automaticity via different biophysical mechanisms involving Ca and membrane clocks, respectively; the clocks are coupled and the intrinsic automaticity is defined by a combined and synergistic action of the clocks as depicted in the *P_up_-g_CaL_* diagram (Maltsev and Lakatta, 2009). Perturbation of ether clock inevitably affects the other clock and the entire coupled-clock system (Yaniv et al., 2013). That is why *g_CaL_* and *P_up_* effectively enhanced robustness of SAN function in our simulations.

More specifically, by allowing *P_up_* and/or *g_CaL_* in the non-firing zone to substantially deviate from their mean values and “penetrate” into the rhythmic firing zone (indicated by double-headed arrows in panels A in Figs 3,4,5), we actually create a sub-population of cells that have intrinsic automaticity in addition to the sub-population of intrinsically non-firing cells (dubbed dormant cells (Kim et al., 2018;Tsutsui et al., 2018;Tsutsui et al., 2021). The two subpopulations of cells intimately interact, determining the ultimate performance (or failure) of the SAN tissue they comprise. Thus, the coupled-clock theory explains, in part, another important result of our study that *g_CaL_* and *P_up_* heterogeneities act synergistically in rescuing SAN from halt. When the SAN system featured heterogeneity in both *g_CaL_* and *P_up_*, almost all cells (~70%, scenario 8) generated APs in marked contrast to SAN tissue with heterogeneity in either *g_CaL_* or *P_up_*, in which all cells remain non-firing (scenarios 4 and 7). Indeed, the respective (purple) double-headed arrow in *P_up_-g_CaL_* diagram in Fig. 5A penetrates deeply into the rhythmic firing zone, enriching SAN tissue with cells featuring high-performance automaticity.

Furthermore, the fact that the SR Ca pumping is a key timing mechanism in the coupled-clock theory can also explain the differences in rescuing tissue automaticity via *P_up_* or *g_CaL_*: AP cycle length substantially varied in tissues with *P_up_* heterogeneity (scenario 5), while only slight AP cycle length variability was observed in tissues with *g_CaL_* heterogeneity (scenarios 1-3). On the other hand, the fact that *I_CaL_* generates AP upstroke, i.e. *g_CaL_* is critical for cell excitability, can explain the presence of non-firing cells in scenarios 1-3. Indeed, as we set *g_CaL_* within a substantial spread (including very low values, like in real SAN cells, Fig. 1A), the entire cell population is comprised of three sub-populations with respect to cell intrinsic properties: (i) spontaneously firing APs; (ii) excitable, but lacking automaticity; (iii) non-excitable. Thus, the intrinsically non-excitable cells do not generate APs in the SAN tissue, explaining our result in scenarios 1-3 involving *g_CaL_* heterogeneity that the percentage of firing cells was always less than 100%. On the other hand, in scenarios involving *P_up_* heterogeneity all or almost all (100% or 98%) cells fired rhythmic APs when *g_CaL_* was chosen relatively high, thereby excluding nonexcitable cells.

### Emergence of SAN tissue automaticity is a critical phenomenon

While the coupled-clock theory has explained, in part, many results of the present study, its application to tissue function is limited, because it predicts only intrinsic cell properties, i.e. cell operation in isolation. Cell tissue, however, operates at the next, higher level of organization and its automaticity emerges from interactions of numerous cells with different intrinsic properties within cell communities. These interactions go beyond the coupled-clock theory (at least in its present form). One important aspect of such emergent behavior was revealed by our simulations: the percentage of firing cells in tissue in all scenarios involving either *P_up_* or *g_CaL_* (except scenarios 4 and 7 lacking automaticity) was much larger than that when the same cells operated in isolation (Table 1), indicating that some excitable cells lacking automaticity were recruited to fire APs by their neighboring cells in local cell communities to increase overall performance of the SAN tissue. And vice versa, the SAN tissue surprisingly lacked automaticity in scenarios 4 or 7, despite their double-headed arrows penetrated deep into the rhythmic firing zone (Fig. 5A), indicating that a major fraction of cells (39% or 30%, respectively, Table 1) was capable of generating rhythmic APs, according to the coupled-clock theory, but did not fire APs.

Thus, our simulations at the edge of the system stability discovered strong amplification (i.e. self-organization) of either AP firing or suppression of AP firing among the cells within SAN tissue. In other terms, whether the system gains automaticity or fails, seems to be a critical phenomenon that depends on intimate interactions of intrinsically firing and non-firing cells (dormant cells). When intrinsically firing cells reach a critical number, they prevail: they not only fire AP themself, but also recruit a fraction of dormant cells to fire APs. And, vice versa, as the sub-population of firing cells decreases, these cells become surrounded by a critical number of non-firing cells, and these non-firing cells suppress the entire system automaticity (for example, in scenarios 4 and 7). At the edge of criticality, the transition from the firing state to non-firing state in SAN tissue is extremely abrupt, like a phase transition in statistical physics. For example, in Movies 4 and 7, the AP generation suddenly ceased at about 3.5 s and 4.5 s after simulation onset, respectively, when the system reached a criticality and the suppressing effect of non-firing cells prevailed.

Because we distributed cells uniformly randomly, the critical phenomenon of the emerging automaticity in our tissue model can be interpreted in terms of Poisson clumping (Aldous, 1989), a phenomenon wherein random events (in space and/or time) have a tendency to occur in clusters, clumps, or bursts. In other words, forming cell clusters with high *g_CaL_* and/or *P_up_* could be a natural consequence of the random distribution. When these natural clusters of intrinsically firing cells are rare and small, their automaticity is suppressed by surrounding dormant cells (which, in turn, also tend to form clusters via Poisson clumping).

### Cell clustering is yet another mechanism to enhance robustness of SAN function

While forming functional clusters via Poisson clumping in our model is a matter of chance (albeit predictable via statistical methods (Aldous, 1989)), the real SAN tissue, as any biological tissue, can direct cell locations to optimize its function. Indeed, high resolution imaging studies in intact SAN detected clusters of cells with various types of functional activity (Fig. 4 in (Bychkov et al., 2020)). Based on this logic, we rescued SAN tissue function in the “hopeless” scenarios 7 and 4 (Movies 4 and 7) just by reorganizing the cell locations within the tissue models (scenarios 9 and 10, Movies 9 and 10). Dormant cells were “kicked” to SAN periphery, while intrinsically firing cells supported AP firing of each other were clustered in the center. As a result, the tissue automaticity was not only rescued, but a substantial fraction of dormant cells (at the border of the firing/non-firing cells) was recruited to fire APs, as evident from the increase of the percentage of firing cells (scenarios 9 and 10 in Table 1). Thus, cell clustering (e.g. with respect to *g_CaL_, P_up_*, or perhaps other parameter) may represent an additional mechanism by which SAN tissue can enhance its robust function at the edge of stability. A large number of local oscillators detected in intact SAN (Bychkov et al., 2020) may reflect cell clustering due to heterogeneity in *I_CaL_* (yielding non-excitable cells, besides excitable cells, as described above, see also Fig. 1A), but the fact that the cell clusters operate at different frequencies point to (among perhaps other interpretations) cell heterogeneity in SR Ca pumping determined by various levels of phospholamban phosphorylation among cells or cell clusters (mimicked by *P_up_* heterogeneity in our model). Indeed, heterogeneity in *P_up_* created numerous oscillators operating at various frequencies in our tissue models (wide histograms in Figs 4B, 5B, 5D).

An apparent downside of cell clustering, however, is that the isolated clusters of automaticity lack physiological importance until their signals overcome the non-firing cell “barriers” to interact with other clusters and/or the rest of the SAN to ultimately deliver synchronized, rhythmic (but not metronomic) APs to atria. While this appears to be a problem in our simple 2-dimensional model of SAN tissue, the real SAN tissue is a 3-dimensional, multilayer meshwork of long and highly branched, intertwined cells (Bychkov et al., 2020). Such tight meshwork of HCN4^+^/Conexin43^-^ cells is structured for signal transfer within and among the functional clusters, regardless of the specific nature of cell interactions that remains unknown.

### Cell heterogeneity is an antiarrhythmic mechanism

Another important result of our study is that heterogeneity in *g_CaL_* or *P_up_* has antiarrhythmic effect. Indeed, while the tissue models (scenarios 3 and 6) with identical cells generated chaotic firing, allowing *g_CaL_* or *P_up_* values to spread (but keeping same averages over the cellular network) shifted the system towards normal, rhythmic operation (Fig. 3D, Movie 3 and Fig. 4C, Movie 6). This result is remarkable and counterintuitive: when the noise of an intrinsically chaotic oscillatory system is combined with the noise of random distribution of its key parameter (*g_CaL_* or *P_up_*) within a SAN network, it is hard to expect that such “double-noisy” system would generate normal rhythmic automaticity; but it does.

### Study limitations and future studies

SAN is an extremely heterogeneous, complex system with respect to individual cell properties, cell interactions, cell network structure, and its autonomic system modulation. The present study tested the hypothesis that heterogeneity of cell functional properties confers robustness to SAN tissue pacemaker function. Our testing focused on two specific key parameters of the cell coupled clock-system, *g_CaL_* and *P_up_* using a simple model of SAN tissue that was a square grid of 25×25 cells of only one cell model type (Maltsev-Lakatta model). Because we study robustness, we further focused our simulations and analysis close to the edge of the system stability and did not examine the entire parametric space of *P_up_ - g_CaL_*. Thus, our study provides evidence (i.e. Proof of Principle) supporting our specific hypothesis and does not include other numerous important factors that determine SAN function. Future numerical studies can explore contributions of other ion currents such as *I_Na_* (Lei et al., 2004), *I_CaT_* (Ono and Iijima, 2005;Tanaka et al., 2008), and funny current (for its heterogeneity data obtained in isolated cells, see (Honjo et al., 1996;Monfredi et al., 2017)), phosphorylation of Ca cycling proteins (phospholamban and RyRs), autonomic modulation, cell network structure, including cell connectivity and interactions in 3 dimensions, interactions with other cell types (e.g. fibroblasts (Camelliti et al., 2004) and telocytes (Mitrofanova et al., 2018)), mechano-sensitivity (Quinn and Kohl, 2012), endogenous Ca buffers, pathological conditions, aging, etc.

## CONCLUSIONS

Natural heterogeneity of pacemaker cells increases robustness of cardiac pacemaker function, i.e. the SAN tissue populated by heterogeneous cells can generate normal automaticity within the cell parameter ranges, whereas SAN populated by identical cells would exhibit either dysrhythmic AP firing or complete lack of automaticity. This effect to enhance pacemaker function at the edge of the system stability is synergistic with respect to heterogeneity of Ca and membrane clocks, and it is not due to a simple summation of activity of intrinsically firing cells that are naturally present in heterogeneous SAN. These firing cells critically interact with intrinsically non-firing cells (dormant cells) and either recruit many of them to fire granting SAN automaticity, or non-firing cells suppress firing cells and SAN tissue automaticity. Clustering cells with specific cell parameters provides an additional mechanism to enhance robustness of SAN function, but can generate isolated communities of firing cells in our simple 2-dimensional model. Therefore, additional mechanisms are required for those clusters to interact with the rest of the SAN. Such mechanisms can be explored in the future in more complex SAN tissue models. Our results provide new insights for SAN operation at the edge of stability that can be helpful not only to understanding normal SAN function especially at low rates and in HCN4^+^/Conexin43^-^ cell communities (Bychkov et al., 2020;Fenske et al., 2020), but also with respect to understanding SAN function in normal aging (linked to deficient cAMP-PKA-Ca signaling (Liu et al., 2014)) and pathological conditions (such as sick sinus syndrome and SAN arrhythmias (Dobrzynski et al., 2007;John and Kumar, 2016)).

## ABBREVIATIONS AND KEY TERMS

SAN: sinoatrial node
*V_m_*: membrane potential
AP: action potential
*I_CaL_*: L-type Ca current
*g_CaL_*: maximum conductance of L-type Ca current
SR: sarcoplasmic reticulum
RyR: ryanodine receptor (Ca release channel)
SERCA2a: SR Ca pump
*P_up_*: maximum rate of SR Ca pumping by SERCA2a
PKA: protein kinase A
CaMKII: Ca/calmodulin-dependent protein kinase II
GPU: graphics processing unit
CUDA: Compute Unified Device Architecture
Non-firing zone: the area of *P_up_–g_CaL_* parametric space where individual SAN cells lack automaticity
Chaotic firing zone: the area of *P_up_–g_CaL_* parametric space where individual SAN cells generate chaotic (dysrhythmic) AP firing
Rhythmic firing zone: the area of *P_up_–g_CaL_* parametric space where individual SAN cells generate rhythmic AP firing
The bifurcation line: the border line in *P_up_–g_CaL_* parametric space separating firing, non-firing, and chaotic firing zones.
Robustness: ability of SAN tissue to generate rhythmic APs in a wide range of model parameters: the larger rhythmic firing zone, the higher robustness of the tissue model.
The coupled-clock system: a coupled system of ion current oscillator in the cell membrane and intracellular Ca oscillator of the SR inside a SAN cell. According to the coupled-clock theory, the oscillators critically interact via multiple Ca- and voltage-dependent mechanisms to generate normal SAN cell automaticity.

## DATA AVAILABILITY STATEMENT

The raw data supporting the conclusions of this article will be made available by the authors, without undue reservation.

## ETHICS STATEMENT

This is a numerical model study. No animal studies are presented in this paper. No human studies are presented in this paper. No potentially identifiable human images or data is presented in this study.

## AUTHOR CONTRIBUTIONS

AVM wrote computer program code, performed model simulations and data analysis. AVM and VAM wrote the first draft of the manuscript. MDS, EGL wrote sections of the manuscript. All authors contributed to conception, design of the study, manuscript revision, read, and approved the submitted version.

## SUPPLEMENTARY MATERIAL

The Supplementary Material for this article can be found online at https://www.frontiersin.org It includes 10 supplementary movies. Each supplementary movie illustrates the result of numerical simulation of SAN tissue (25×25 cells) lasted 7.5 s. in the respective model scenario described in Table 1. The model parameters in each movie are also summarized below. *V_m_* dynamics in each cell in the grid was color-coded via red shade from −60 mV (pure black) to 20 mV (pure red). Simulation time is displayed in the upper left corner.

**Supplementary Movie 1.** *P_up_* = 7 mM/s, *g_CaL_* ∈ [0.14, 0.54] nS/pF, mean = 0.34 nS/pF

**Supplementary Movie 2.** *P_up_* = 5 mM/s, *g_CaL_* ∈ [0.17, 0.57] nS/pF, mean = 0.37 nS/pF

**Supplementary Movie 3.** *P_up_* = 3 mM/s, *g_CaL_* ∈ [0.21, 0.61] nS/pF, mean = 0.41 nS/pF

**Supplementary Movie 4.** *P_up_* ∈ [2, 12], mean = 7 mM/s, *g_CaL_* = 0.34 nS/pF

**Supplementary Movie 5.** *P_up_* ∈ [0, 10], mean = 5 mM/s, *g_CaL_* = 0.37 nS/pF

**Supplementary Movie 6.** *P_up_* ∈ [0, 6], mean = 3 mM/s, *g_CaL_* = 0.41 nS/pF

**Supplementary Movie 7.** *P_up_* = 7 mM/s, *g_CaL_* ∈ [0.3, 0.38] nS/pF, mean = 0.34 nS/pF

**Supplementary Movie 8.** *P_up_* ∈ [2, 12], mean =75 mM/s, *g_CaL_* ∈ [0.3, 0.38] nS/pF, mean = 0.34 nS/pF

**Supplementary Movie 9.** *P_up_* = 7 mM/s, g_CaL_ ∈ [0.3, 0.38] nS/pF, mean = 0.34 nS/pF, cells with higher *g_CaL_* were clustered to SAN center

**Supplementary Movie 10.** *P_up_* ∈ [2, 12], mean = 7 mM/s with *g_CaL_* = 0.34 nS/pF, cells with higher *P_up_* were clustered to SAN center

## REFERENCES

Aldous, D. (1989). Probability Approximations via the Poisson Clumping Heuristic. Applied Mathematical Sciences 7, Springer (book).

Bleeker, W.K., Mackaay, A.J., Masson-Pevet, M., Bouman, L.N., and Becker, A.E. (1980). Functional and morphological organization of the rabbit sinus node. Circ Res 46, 11–22.

Boyett, M.R., Honjo, H., and Kodama, I. (2000). The sinoatrial node, a heterogeneous pacemaker structure. Cardiovasc Res 47, 658–687.

Bychkov, R., Juhaszova, M., Tsutsui, K., Coletta, C., Stern, M.D., Maltsev, V.A., and Lakatta, E.G. (2020). Synchronized cardiac impulses emerge from multi-scale, heterogeneous local calcium signals within and among cells of heart pacemaker tissue. JACC Clin Electrophysiol 6, 907–931. doi: 10.1016/j.jacep.2020.06.022.

Camelliti, P., Green, C.R., Legrice, I., and Kohl, P. (2004). Fibroblast network in rabbit sinoatrial node: structural and functional identification of homogeneous and heterogeneous cell coupling. Circ Res 94, 828–835. doi: 10.1161/01.RES.0000122382.19400.14.

Campana, C.A. (2015). 2-Dimensional Computational Model to Analyze the Effects of Cellular Heterogeinity on Cardiac Pacemaking. PhD thesis, Bologna, Italy: Universita di Bologna, Corso di Studio in Ingegneria Biomedical.

Clancy, C.E., and Santana, L.F. (2020). Evolving Discovery of the Origin of the Heartbeat: A New Perspective on Sinus Rhythm. JACC Clin Electrophysiol 6, 932–934. doi: 10.1016/j.jacep.2020.07.002.

Dobrzynski, H., Boyett, M.R., and Anderson, R.H. (2007). New insights into pacemaker activity: promoting understanding of sick sinus syndrome. Circulation 115, 1921–1932.

Fenske, S., Hennis, K., Rotzer, R.D., Brox, V.F., Becirovic, E., Scharr, A., Gruner, C., Ziegler, T., Mehlfeld, V., Brennan, J., Efimov, I.R., Pauza, A.G., Moser, M., Wotjak, C.T., Kupatt, C., Gonner, R., Zhang, R., Zhang, H., Zong, X., Biel, M., and Wahl-Schott, C. (2020). cAMP-dependent regulation of HCN4 controls the tonic entrainment process in sinoatrial node pacemaker cells. Nat Commun 11, 5555. doi: 10.1038/s41467-020-19304-9.

Garny, A., Noble, D., Hunter, P.J., and Kohl, P. (2009). CELLULAR OPEN RESOURCE (COR): current status and future directions. Philos Trans A Math Phys Eng Sci 367, 1885–1905. doi: 10.1098/rsta.2008.0289.

Glynn, P., Onal, B., and Hund, T.J. (2014). Cycle length restitution in sinoatrial node cells: a theory for understanding spontaneous action potential dynamics. PLoS One 9, e89049. doi: 10.1371/journal.pone.0089049.

Gratz, D., Onal, B., Dalic, A., and Hund, T.J. (2018). Synchronization of pacemaking in the sinoatrial node: a mathematical modeling study. Front Physics 6, id.63. doi: doi.org/10.3389/fphy.2018.00063.

Honjo, H., Boyett, M.R., Kodama, I., and Toyama, J. (1996). Correlation between electrical activity and the size of rabbit sino-atrial node cells. J Physiol 496 ( Pt 3), 795–808.

Honjo, H., Lei, M., Boyett, M.R., and Kodama, I. (1999). Heterogeneity of 4-aminopyridine-sensitive current in rabbit sinoatrial node cells. Am J Physiol 276, H1295–1304.

Huang, X., Mi, Y., Qian, Y., and Hu, G. (2011). Phase-locking behaviors in an ionic model of sinoatrial node cell and tissue. Phys Rev E Stat Nonlin Soft Matter Phys 83, 061917. doi: 10.1103/PhysRevE.83.061917.

Imtiaz, M.S., Von Der Weid, P.Y., Laver, D.R., and Van Helden, D.F. (2010). SR Ca^2+^ store refill--a key factor in cardiac pacemaking. J Mol Cell Cardiol 49, 412–426.

Jalife, J. (1984). Mutual entrainment and electrical coupling as mechanisms for synchronous firing of rabbit sino-atrial pace-maker cells. J Physiol 356, 221–243.

John, R.M., and Kumar, S. (2016). Sinus Node and Atrial Arrhythmias. Circulation 133, 1892–1900. doi: 10.1161/CIRCULATIONAHA.116.018011.

Karpaev, A.A., Syunyaev, R.A., and Aliev, R.R. (2018). Effects of fibroblast-myocyte coupling on the sinoatrial node activity: A computational study. Int J Numer Method Biomed Eng 34, e2966. doi: 10.1002/cnm.2966.

Keith, A., and Flack, M. (1907). The Form and Nature of the Muscular Connections between the Primary Divisions of the Vertebrate Heart. J Anat Physiol 41, 172–189.

Kharche, S.R., Vigmond, E., Efimov, I.R., and Dobrzynski, H. (2017). Computational assessment of the functional role of sinoatrial node exit pathways in the human heart. PLoS One 12, e0183727. doi: 10.1371/journal.pone.0183727.

Kim, M.S., Maltsev, A.V., Monfredi, O., Maltseva, L.A., Wirth, A., Florio, M.C., Tsutsui, K., Riordon, D.R., Parsons, S.P., Tagirova, S., Ziman, B.D., Stern, M.D., Lakatta, E.G., and Maltsev, V.A. (2018). Heterogeneity of calcium clock functions in dormant, dysrhythmically and rhythmically firing single pacemaker cells isolated from SA node. Cell Calcium 74, 168–179. doi: 10.1016/j.ceca.2018.07.002.

Kim, M.S., Monfredi, O., Maltseva, L.A., Lakatta, E.G., and Maltsev, V.A. (2021). beta-Adrenergic Stimulation Synchronizes a Broad Spectrum of Action Potential Firing Rates of Cardiac Pacemaker Cells toward a Higher Population Average. Cells 10. doi: 10.3390/cells10082124.

Kurata, Y., Hisatome, I., Imanishi, S., and Shibamoto, T. (2002). Dynamical description of sinoatrial node pacemaking: improved mathematical model for primary pacemaker cell. Am J Physiol 283, H2074–2101.

Kurata, Y., Hisatome, I., and Shibamoto, T. (2012). Roles of Sarcoplasmic Reticulum Ca^2+^Cycling and Na^+^/Ca^2+^ Exchanger in Sinoatrial Node Pacemaking: insights from bifurcation analysis of mathematical models. Am J Physiol Heart Circ Physiol 302, H2285–H2300.

Lakatta, E.G., Maltsev, V.A., and Vinogradova, T.M. (2010). A coupled SYSTEM of intracellular Ca^2+^ clocks and surface membrane voltage clocks controls the timekeeping mechanism of the heart’s pacemaker. Circ Res 106, 659–673.

Lei, M., Honjo, H., Kodama, I., and Boyett, M.R. (2001). Heterogeneous expression of the delayed-rectifier K+ currents i(K,r) and i(K,s) in rabbit sinoatrial node cells. J Physiol 535, 703–714.

Lei, M., Jones, S.A., Liu, J., Lancaster, M.K., Fung, S.S., Dobrzynski, H., Camelliti, P., Maier, S.K., Noble, D., and Boyett, M.R. (2004). Requirement of neuronal- and cardiac-type sodium channels for murine sinoatrial node pacemaking. J Physiol 559, 835–848.

Li, J., Inada, S., Schneider, J.E., Zhang, H., Dobrzynski, H., and Boyett, M.R. (2014). Three-dimensional computer model of the right atrium including the sinoatrial and atrioventricular nodes predicts classical nodal behaviours. PLoS One 9, e112547. doi: 10.1371/journal.pone.0112547.

Li, K., Chu, Z., and Huang, X. (2018). Annihilation of the pacemaking activity in the sinoatrial node cell and tissue AIP Advances 8, 125319.

Li, P., Lines, G., Maleckar, M., and Tveito, A. (2013). Mathematical models of cardiac pacemaking function. Frontiers in Physics 1. doi: 10.3389/fphy.2013.00020.

Liu, J., Sirenko, S., Juhaszova, M., Sollott, S.J., Shukla, S., Yaniv, Y., and Lakatta, E.G. (2014). Age-associated abnormalities of intrinsic automaticity of sinoatrial nodal cells are linked to deficient cAMP-PKA-Ca^2+^ signaling. Am J Physiol Heart Circ Physiol 306, H1385–1397. doi: 10.1152/ajpheart.00088.2014.

Luo, C.H., and Rudy, Y. (1994). A dynamic model of the cardiac ventricular action potential. I. Simulations of ionic currents and concentration changes. Circ Res 74, 1071–1096.

Lyashkov, A.E., Behar, J., Lakatta, E.G., Yaniv, Y., and Maltsev, V.A. (2018). Positive Feedback Mechanisms among Local Ca Releases, NCX, and ICaL Ignite Pacemaker Action Potentials. Biophys J 114, 1176–1189. doi: 10.1016/j.bpj.2017.12.043.

Maltsev, A.V., Maltsev, V.A., and Stern, M.D. (2017). Stabilization of diastolic calcium signal via calcium pump regulation of complex local calcium releases and transient decay in a computational model of cardiac pacemaker cell with individual release channels. PLoS Comput Biol 13,e1005675. doi: 10.1371/journal.pcbi.1005675.

Maltsev, V.A., and Lakatta, E.G. (2009). Synergism of coupled subsarcolemmal Ca^2+^ clocks and sarcolemmal voltage clocks confers robust and flexible pacemaker function in a novel pacemaker cell model. Am J Physiol Heart Circ Physiol 296, H594–H615.

Maltsev, V.A., and Lakatta, E.G. (2010). A novel quantitative explanation for autonomic modulation of cardiac pacemaker cell automaticity via a dynamic system of sarcolemmal and intracellular proteins. Am J Physiol Heart Circ Physiol 298, H2010–H2023.

Maltsev, V.A., and Lakatta, E.G. (2013). Numerical models based on a minimal set of sarcolemmal electrogenic proteins and an intracellular Ca clock generate robust, flexible, and energy-efficient cardiac pacemaking. J Mol Cell Cardiol 59, 181–195.

Maltsev, V.A., Yaniv, Y., Maltsev, A.V., Stern, M.D., and Lakatta, E.G. (2014). Modern perspectives on numerical modeling of cardiac pacemaker cell. J Pharmacol Sci 125, 6–38.

Mangoni, M.E., and Nargeot, J. (2008). Genesis and regulation of the heart automaticity. Physiol Rev 88, 919–982.

Mata, A.N., Alonso, G.R., Garza, G.L., Fernández, J.R.G., García, M.a.C., and Ábrego, N.P.C. (2019). Parallel simulation of the synchronization of heterogeneous cells in the sinoatrial node. Concurrency Computat Pract Exper, e5317. doi: DOI: 10.1002/cpe.5317.

Mattick, P., Parrington, J., Odia, E., Simpson, A., Collins, T., and Terrar, D. (2007). Ca^2+^-stimulated adenylyl cyclase isoform AC1 is preferentially expressed in guinea-pig sinoatrial node cells and modulates the I_f_ pacemaker current. J Physiol 582, 1195–1203.

Michaels, D.C., Matyas, E.P., and Jalife, J. (1987). Mechanisms of sinoatrial pacemaker synchronization: a new hypothesis. Circ Res 61, 704–714.

Mitrofanova, L.B., Gorshkov, A.N., Konovalov, P.V., and Krylova, J.S. (2018). Telocytes in the human sinoatrial node. J Cell Mol Med 22, 521–532. doi: 10.1111/jcmm.13340.

Monfredi, O., Tsutsui, K., Ziman, B., Stern, M.D., Lakatta, E.G., and Maltsev, V.A. (2018). Electrophysiological heterogeneity of pacemaker cells in the rabbit intercaval region, including the SA node: insights from recording multiple ion currents in each cell. Am J Physiol Heart Circ Physiol 314, H403–H414. doi: 10.1152/ajpheart.00253.2016.

Monfredi, O., Tsutsui, K., Ziman, B.D., Stern, M.D., Lakatta, E.G., and Maltsev, V.A. (2017). Electrophysiological heterogeneity of pacemaker cells in rabbit intercaval region, including SA node: insights from recording multiple ion currents in each cell. Am J Physiol Heart Circ Physiol, ajpheart 00253 02016. doi: 10.1152/ajpheart.00253.2016.

Munoz, M.A., Kaur, J., and Vigmond, E.J. (2011). Onset of atrial arrhythmias elicited by autonomic modulation of rabbit sinoatrial node activity: a modeling study. Am J Physiol Heart Circ Physiol 301, H1974–1983. doi: 10.1152/ajpheart.00059.2011.

Musa, H., Lei, M., Honjo, H., Jones, S.A., Dobrzynski, H., Lancaster, M.K., Takagishi, Y., Henderson, Z., Kodama, I., and Boyett, M.R. (2002). Heterogeneous expression of Ca^2+^handling proteins in rabbit sinoatrial node. J Histochem Cytochem 50, 311–324.

Ono, K., and Iijima, T. (2005). Pathophysiological significance of T-type Ca^2+^ channels: properties and functional roles of T-type Ca^2+^ channels in cardiac pacemaking. J Pharmacol Sci 99, 197–204.

Oren, R.V., and Clancy, C.E. (2010). Determinants of heterogeneity, excitation and conduction in the sinoatrial node: a model study. PLoS Comput Biol 6, e1001041.

Quinn, T.A., and Kohl, P. (2012). Mechano-sensitivity of cardiac pacemaker function: pathophysiological relevance, experimental implications, and conceptual integration with other mechanisms of rhythmicity. Prog Biophys Mol Biol 110, 257–268. doi: 10.1016/j.pbiomolbio.2012.08.008.

Sano, T., Sawanobori, T., and Adaniya, H. (1978). Mechanism of rhythm determination among pacemaker cells of the mammalian sinus node. Am J Physiol 235, H379–384. doi: 10.1152/ajpheart.1978.235.4.H379.

Severi, S., Fantini, M., Charawi, L.A., and Difrancesco, D. (2012). An updated computational model of rabbit sinoatrial action potential to investigate the mechanisms of heart rate modulation. J Physiol 590, 4483–4499.

Sirenko, S.G., Maltsev, V.A., Yaniv, Y., Bychkov, R., Yaeger, D., Vinogradova, T., Spurgeon, H.A., and Lakatta, E.G. (2016). Electrochemical Na^+^ and Ca^2+^ gradients drive coupled-clock regulation of automaticity of isolated rabbit sinoatrial nodal pacemaker cells. Am J Physiol Heart Circ Physiol 311, H251–267. doi: 10.1152/ajpheart.00667.2015.

Stern, M.D., Maltseva, L.A., Juhaszova, M., Sollott, S.J., Lakatta, E.G., and Maltsev, V.A. (2014). Hierarchical clustering of ryanodine receptors enables emergence of a calcium clock in sinoatrial node cells. J Gen Physiol 143, 577–604.

Tanaka, H., Komikado, C., Namekata, I., Nakamura, H., Suzuki, M., Tsuneoka, Y., Shigenobu, K., and Takahara, A. (2008). Species difference in the contribution of T-type calcium current to cardiac pacemaking as revealed by r(-)-efonidipine. J Pharmacol Sci 107, 99–102.

Torrente, A.G., Mesirca, P., Neco, P., Rizzetto, R., Dubel, S., Barrere, C., Sinegger-Brauns, M., Striessnig, J., Richard, S., Nargeot, J., Gomez, A.M., and Mangoni, M.E. (2016). L-type Cav1.3 channels regulate ryanodine receptor-dependent Ca^2+^ release during sino-atrial node pacemaker activity. Cardiovasc Res 109, 451–461. doi: 10.1093/cvr/cvw006.

Tsutsui, K., Florio, M.C., Yang, A., Wirth, A.N., Yang, D., Kim, M.S., Ziman, B.D., Bychkov, R., Monfredi, O.J., Maltsev, V.A., and Lakatta, E.G. (2021). cAMP-Dependent Signaling Restores AP Firing in Dormant SA Node Cells via Enhancement of Surface Membrane Currents and Calcium Coupling. Front Physiol 12, 596832. doi: 10.3389/fphys.2021.596832.

Tsutsui, K., Monfredi, O., Sirenko-Tagirova, S.G., Maltseva, L.A., Bychkov, R., Kim, M.S., Ziman, B.D., Tarasov, K.V., Tarasova, Y.S., Zhang, J., Wang, M., Maltsev, A.V., Brennan, J.A., Efimov, I.R., Stern, M.D., Maltsev, V.A., and Lakatta, E.G. (2018). A coupled-clock system drives the automaticity of human sinoatrial nodal pacemaker cells. Sci Signal 11, eaap7608.

Vinogradova, T.M., Brochet, D.X., Sirenko, S., Li, Y., Spurgeon, H., and Lakatta, E.G. (2010). Sarcoplasmic reticulum Ca^2+^ pumping kinetics regulates timing of local Ca^2+^ releases and spontaneous beating rate of rabbit sinoatrial node pacemaker cells. Circ Res 107, 767–775.

Vinogradova, T.M., and Lakatta, E.G. (2009). Regulation of basal and reserve cardiac pacemaker function by interactions of cAMP mediated PKA-dependent Ca^2+^ cycling with surface membrane channels. Journal of Molecular and Cellular Cardiology 47, 456–474.

Weiss, J.N., and Qu, Z. (2020). The Sinus Node: Still Mysterious After All These Years. JACC Clin Electrophysiol 6, 1841–1843. doi: 10.1016/j.jacep.2020.09.017.

Yang, D., Morrell, C.H., Lyashkov, A.E., Tagirova, S., Zahanich, I., Yaniv, Y., Vinogradova, T., Ziman, B., Maltsev, V.A., and Lakatta, E.G. (2021). Ca2+ and Membrane Potential Transitions During Action Potentials are Self-Similar to Each Other and to Variability of AP Firing Intervals Across the Broad Physiologic Range of AP Intervals During Autonomic Receptor Stimulation. Front Physiol in print. doi: DOI: 10.3389/fphys.2021.612770.

Yaniv, Y., Sirenko, S., Ziman, B.D., Spurgeon, H.A., Maltsev, V.A., and Lakatta, E.G. (2013). New evidence for coupled clock regulation of the normal automaticity of sinoatrial nodal pacemaker cells: Bradycardic effects of ivabradine are linked to suppression of intracellular Ca cycling. J Mol Cell Cardiol 62C, 80–89.

Younes, A., Lyashkov, A.E., Graham, D., Sheydina, A., Volkova, M.V., Mitsak, M., Vinogradova, T.M., Lukyanenko, Y.O., Li, Y., Ruknudin, A.M., Boheler, K.R., Van Eyk, J., and Lakatta, E.G. (2008). Ca^2+^-stimulated basal adenylyl cyclase activity localization in membrane lipid microdomains of cardiac sinoatrial nodal pacemaker cells. J Biol Chem 283, 14461–14468.

Zhang, H., Holden, A.V., and Boyett, M.R. (2001). Gradient model versus mosaic model of the sinoatrial node. Circulation 103, 584–588.

Zhang, Y., Ocampo-Espindola, J.L., Kiss, I.Z., and Motter, A.E. (2021). Random heterogeneity outperforms design in network synchronization. Proc Natl Acad Sci U S A 118. doi: 10.1073/pnas.2024299118.

